# Tunable Chemical and Optical Control of ER-Plasma Membrane Contact Site Geometry and Dynamics with High-Fidelity Visualization

**DOI:** 10.64898/2026.01.28.701813

**Authors:** Shaoqing Zhang, Jinyu Fei, Yuanmin Zheng, Ruobo Zhou

## Abstract

Endoplasmic reticulum-plasma membrane (ER-PM) contact sites are essential signaling hubs that regulate lipid transport, calcium homeostasis, and spatially organized signal transduction. Emerging evidence indicates that not only the presence but also the dynamics, stability, and geometry of ER-PM contacts critically shape cellular functions; however, tools that enable simultaneous high-fidelity visualization and reversible, quantitative control of these contacts in living cells remain limited. Here, we introduce a modular toolkit for inducible ER-PM contact-site reconstitution based on complementary chemical and optical dimerization strategies. We develop a nontoxic and reversible abscisic acid (ABA)-inducible system using the plant-derived ABIcs/PYLcs pair, and a rapidly reversible optogenetic system based on the iLID/SspB module, both of which allow robust visualization and dose-dependent control over contact-site formation kinetics, increasing contact-site density and total area fraction per cell without altering the size of individual contacts. In contrast, systematic variation of rigid α-helical linker length or inducible tether abundance selectively tunes the lateral growth, stability, and lifetime of individual contact sites, without changing their density. By combining these two orthogonal strategies, we achieve independent control of both individual contact-site size and overall contact-site density, providing complementary mechanisms to adjust total contact area per cell. This versatile platform enables quantitative dissection of ER-PM contact site structure-function relationships and offers broad utility in studies of lipid exchange, calcium signaling, membrane repair, metabolic regulation, and disease-relevant dysregulation.

## INTRODUCTION

Membrane contact sites (MCSs) are specialized zones where two membrane-bound cellular organelles are positions in close apposition without undergoing fusion, maintained by dedicated protein- and lipid-based tethers^1^. MCSs support a broad array of essential cellular functions, including non-vesicular lipid transport, Ca² and ion homeostasis, organelle positioning and dynamics, metabolic coordination, signaling compartmentalization, autophagy regulation, and cellular stress responses^2^. Among the diverse classes of MCSs, endoplasmic reticulum-plasma membrane (ER-PM) contact sites constitute one of the most abundant and functionally versatile MCS types in mammalian cells^3^. ER-PM contacts form a widespread (*i.e.*, covers 2-5% of the PM surface in many cell types) yet highly regulated subcellular interface that supports lipid transfer, Ca² signaling and homeostasis, phosphoinositide metabolism, cell signaling polarity, cell migration, and membrane repair^4–6^. Dysregulation of ER-PM contact sites has been implicated in numerous pathological conditions, including neurodegenerative disorders^7,8^, muscular hypotonia and related myopathies^9^, immune deficiencies^10^, and cardiovascular and metabolic abnormalities^11^. A well-characterized example of ER-PM function is Ca² signaling: depletion of ER Ca² activates the ER-resident Ca² sensor Stromal Interaction Molecule 1 (STIM1), which translocates to ER-PM junctions and engages the PM-resident Orai1 Ca² channel to drive store-operated calcium entry (SOCE)^12,13^. ER-PM contacts thereby regulate Orai1-dependent SOCE to support muscle growth and reduce fatigue in adult skeletal muscle^14^. In addition, beyond their mere formation, ER-PM contact sites are increasingly recognized as tunable physical interfaces whose nanoscale geometry and dynamics encode functional specificity^15–17^. Native ER-PM contacts span a broad range of intermembrane distances, sizes, and lifetimes, and these parameters vary across cell types, subcellular regions, and physiological states. Accumulating evidence suggests that such geometric heterogeneity is not incidental but functionally consequential, influencing local lipid exchange, calcium signaling, cytoskeletal coupling, and membrane mechanics. For instant migrating cells, ER-PM contact sites exhibit a pronounced front-back gradient in both density and size, with small, transient contacts at the leading edge and large, stable contacts at the trailing edge. This spatial organization of ER-PM contacts is a key determinant of directional signaling and cell migration^15^. Given their indispensable roles in lipid exchange, Ca² dynamics, and cellular signaling, precise tools for visualizing and acutely manipulating ER-PM contact sites in living cells are critically needed. Such systems are essential for mechanistically dissecting their contributions to normal physiology and understanding how their dysregulation drives diseases.

Although several molecular tools have been developed for either visualizing or manipulating ER-PM contact sites, none allow simultaneous high-fidelity visualization and reversible control of these junctions. Genetically encoded reporter MAPPER^18^ (“membrane-attached peripheral ER”) fuses an ER-targeting sequence with a linker and a phosphoinositide-binding motif from the Rit polybasic (PB) domain to associate with the PM. While useful for labeling ER-PM contacts, MAPPER cannot manipulate the dynamic formation and dissociation of ER-PM contacts, and its constitutive tethering may perturb endogenous phosphoinositide homeostasis^19,20^. Optogenetic approaches, such as LiMETER, are based on the MAPPER design and incorporate the AsLOV2 light-sensing domain upstream of the Rit-PB domain^21^. In the dark, AsLOV2 inhibits Rit-PB binding to the plasma membrane, whereas blue light illumination triggers Rit-PB-PM interaction, enabling reversible, noninvasive, and precise manipulation of ER-PM contacts. However, because the LiMETER construct is tagged with fluorescent proteins and anchored to the dense reticular ER network, ER-PM contacts remain partially obscured within the ER mesh. As a result, even after light-induced formation of ER-PM sites, accurate visualization and quantitative analysis are challenging, despite successful acute and reversible control of contact formation. Chemically inducible dimerization (CID) systems provide an alternative strategy to control ER-PM contact site formation with small-molecule inducers. The rapamycin-based FRB-FKBP system has been used to tether the ER and PM, demonstrating that ER-PM contacts serve as platforms for STIM1-Orai1 interactions during store-operated Ca² entry^22^. Despite its utility in chemically inducing heterodimerization between FKBP-tagged and FRB-tagged proteins^23–25^, rapamycin-based CID suffers from extremely slow dissociation kinetics^26^, interference with endogenous mTOR signaling^27^, potential cytotoxicity^28,29^, high cost, and reduced specificity due to competition from endogenous FKBP or cyclophilin family proteins in eukaryote cells^30^. Together, these limitations highlight the need for tools that enable both precise, reversible manipulation and high-fidelity visualization of ER-PM contact sites, enabling quantitative investigation of their dynamic contributions to normal cellular function and pathological processes.

Here, we present a modular and quantitative toolkit that enables high-fidelity visualization and reversible, tunable control of ER-PM contact sites in living cells, addressing key limitations of existing approaches. We developed two complementary inducible tethering platforms: a chemically inducible system based on the plant-derived abscisic acid (ABA)-dependent ABIcs/PYLcs interaction^31^, and an optogenetic system based on the blue-light-responsive iLID/SspB pair^32^. Both systems allow reversible assembly of ER-PM contact sites while providing a spatially confined fluorescent readout for accurate visualization and quantitative analysis. The ABA-based system offers nontoxic, specific, and reversible chemical control without interference from endogenous mammalian signaling, whereas the iLID/SspB system provides fast, noninvasive, reversible optical control with high temporal precision. Beyond inducible formation, we introduce orthogonal, design-driven strategies to modulate contact-site architecture: tuning ABA concentration or blue-light intensity primarily controls contact-site density and total cellular contact area without altering the size of individual contacts, whereas systematic variation of rigid α-helical linker length or inducible control of tether abundance selectively modulates the size and stability of individual contacts without affecting their density. Together, these approaches provide independent and quantitative control over contact-site dynamics, geometry, and extent, offering a versatile framework for dissecting how ER-PM organization influences lipid transport, calcium signaling, membrane repair, metabolic coordination, and broader cellular functions.

## EXPERIMENTAL SECTION

### Plasmid construction

All plasmids and DNA fragments used in this study were obtained from Addgene or synthesized by commercial providers. Coding sequences were amplified by PCR using high-fidelity DNA polymerase (New England Biolabs, M0491L), and plasmids were assembled using Gibson assembly (New England Biolabs, E5510S) according to the manufacturer’s instructions. Unless otherwise noted, constructs were cloned into mammalian expression vectors under the control of a CMV promoter.

For the ER-PM contact-forming chemically induced dimerization (CID) and light-induced dimerization (LID) systems, rapamycin-inducible (FBR-mRFP-T2A-FKBP), abscisic acid (ABA)-inducible (ABIcs-mRFP-T2A-PYLcs), and blue-light inducible (iLID-mRFP-T2A-SspB) expression cassettes were generated. Individual coding sequences were derived from Addgene plasmids (#108137 and #40896; rapamycin system, #134999; ABA system, and #171038; LID system), amplified by PCR, and assembled into a single open reading frame containing a self-cleaving T2A peptide between the two fusion proteins. For PM-targeted constructs, the N-terminal palmitoylation/myristoylation signal of Lyn kinase (MGCIKSKGKDS) was fused in-frame to the N terminus of FRB (rapamycin system), ABIcs (ABA system), or iLID (LID system), via a flexible linker (AGADPTRSANSGAGAGAGAILSR). For ER-targeted constructs, the C-terminal ER localization sequence of human Sac1 phosphatase (residues 521-587) was fused in-frame to the C terminus of FKBP (rapamycin system), PYLcs (ABA system), or SspB (LID system) via a flexible linker (GSGAGAGAGAILNSRV). In all constructs, the PM-targeted and ER-targeted fusion proteins were encoded within a single bicistronic expression cassette and separated by a T2A peptide, enabling stoichiometric expression of the two proteins from a single promoter through ribosomal skipping. To generate variants with altered linker rigidity and length, synthetic DNA fragments encoding α-helical linkers ((EAAAR)_4_ or (EAAAR)_9_; Twist Bioscience) were inserted to the original short linker (*i.e.*, (EAAAR)_0_) by PCR amplification and Gibson assembly.

For XLone-ABA and XLone-iLID/SspB constructs, the constitutive CMV-driven expression cassette was subcloned into the XLone backbone (Addgene, plasmid# 96930) using PCR amplification and Gibson assembly. All plasmids were verified by Sanger DNA sequencing.

### Cell culture

Human osteosarcoma (U2OS) and human embryonic kidney 293T (HEK293T) cells were maintained in Dulbecco’s Modified Eagle Medium (DMEM; Corning, 10-013-CV) supplemented with 10% (v/v) fetal bovine serum (FBS; Gibco, A52567-01) and 1% (v/v) penicillin-streptomycin (Gibco, 10378-016). Cells were maintained at 37 °C in a humidified incubator with 5% CO. For routine passaging, cells were dissociated using 0.25% trypsin-EDTA (Corning, 25-053-CI). For imaging experiments, cells were seeded onto 18-mm round glass coverslips (Electron Microscopy Sciences, 72222-01) placed in 12-well tissue culture plates.

### Cell transfection

Plasmid transfection was performed using TransIT®-LT1 reagent (Mirus Bio, MIR2300) according to the manufacturer’s instructions. Briefly, cells at 60-80% confluency were transfected with 1.0 µg of plasmid DNA diluted in 100 µL of Opti-MEM (Gibco, 31985-070). Subsequently, 3 µL of TransIT-LT1 reagent was added to the diluted DNA, gently mixed, and incubated for 15 min at room temperature (RT) to allow DNA-lipid complex formation. The transfection mixture was then added dropwise to cells cultured in complete growth medium. After 6-12 hours, the medium was replaced with fresh DMEM supplemented with 10% FBS to facilitate cell recovery and transgene expression. For experiments using Xlone-based inducible constructs, doxycycline (Sigma, D3447-500mg) at the indicated concentrations was added to the culture medium 48 h after transfection to induce transgene expression. Cells were subjected to live-cell imaging or fixed for downstream analyses 24-48 h post-transfection, depending on the experimental design.

### ABA and rapamycin induction and withdrawal assays

For live-cell imaging experiments, 100 mM ABA (TCI America, A1698) and 100 mM rapamycin (MilliporeSigma, 553210) stock solutions were diluted to the indicated working concentrations in Gibco™ Live Cell Imaging Solution (LCIS; Thermo Fisher Scientific, A14291DJ). U2OS cells were pre-equilibrated by rinsing once with pre-warmed imaging solution and incubating at 37 °C for at least 15 minutes prior to stimulation. ABA or rapamycin was then added directly to the imaging chamber during image acquisition, and confocal imaging was performed over a 0-30 minute time course following compound addition to monitor ER-PM contact site formation. For withdrawal experiments, cells were treated with ABA or rapamycin for 30 minutes, after which the compounds were removed by washing the cells four times with pre-warmed LCIS. The cells were then maintained in fresh LCIS, and confocal imaging was performed over a 0-7 hour time course following compound withdrawal to assess the reversibility of ER-PM contact site formation.

For fixed-cell assays, ABA stock solutions (100 mM in DMSO) were aliquoted and stored at -20 °C. Before fixation, cells were treated with 100 μM ABA diluted in complete DMEM (with 10% FBS, 1% penicillin-streptomycin) and incubated at 37 °C for 30 minutes. Following induction, cells were immediately washed with phosphate-buffered saline (PBS) and fixed using 4% (w/v) paraformaldehyde containing 4% (w/v) sucrose in PBS for downstream procedures.

### Blue light induction and withdrawal assay

For live-cell imaging assays, U2OS cells were pre-equilibrated by rinsing once with pre-warmed LCIS and incubating at 37 °C for at least 15 minutes prior to light stimulation. Blue light illumination was applied using a 470-nm laser line (89 North) at 5-40% output power, corresponding to a laser power of 0.022-0.177 mW measured at the sample plane using an optical power meter (ThorLabs, PM130D). Confocal imaging was performed during continuous blue light illumination over a 0-15 minute time course to monitor ER-PM contact site formation. For withdrawal experiments, blue light illumination was terminated after 15 minutes of stimulation. Cells were then maintained in fresh LCIS without blue light exposure, and confocal imaging was performed over a 0-15 minute time course following blue light withdrawal to assess the reversibility of ER-PM contact site formation.

For fixed-cell assays, HEK293T cells expressing XLone-based constructs were stimulated prior to fixation using a custom-built blue light illumination system. The system consisted of a 465-nm blue LED light source powered by a DC power supply, with output light intensity adjusted to the desired level. Cells were exposed to continuous blue light illumination for defined durations. To minimize unintended activation, cells were shielded from ambient blue light prior to stimulation. Immediately following stimulation, cells were fixed using 4% (w/v) paraformaldehyde containing 4% (w/v) sucrose in PBS for downstream procedures.

### Widefield epi-fluorescence imaging

Widefield epi-fluorescence imaging was performed using a custom-built microscope based on an Olympus IX-73 body, equipped with a 20× (N.A. 0.80) UPlanXApo air-immersion objective (Olympus), a 60× (N.A. 1.4) PlanApo oil-immersion objective (Olympus), and a CMOS Camera (Thorlabs, CS126MU). A Lumencor SOLA light engine (SOLA U-nIR) was used as the light source for illumination. Unless otherwise indicated, the 60× oil-immersion objective was used to image U2OS cells during rapamycin- and ABA-induced time-course and recovery experiments.

### Confocal imaging

Confocal imaging was performed using a custom-built microscope based on an Olympus IX-83 inverted microscope body, equipped with a CrestOptics X-Light V3 spinning-disk confocal unit. Illumination was provided by an LDI-NIR-7 Laser Diode Illuminator (89 North). Images were collected using either a 60× (NA 1.42) UPlanXApo or a 100× (NA 1.40) UPlanSApo oil-immersion objective (Olympus) and recorded with an ORCA-Fusion BT digital CMOS camera (Hamamatsu, C15440-20UP). Image acquisition was controlled through the Multi-Dimensional Acquisition module in Micro-Manager 2.0.0, enabling multichannel and z-stack imaging. Z- series were acquired from the apical to basal planes of U2OS cells with a step size of 0.2-0.5 µm. Final confocal images were generated by maximum-intensity projection of the z-stacks.

### MTT cell viability assay

Cell viability was assessed using the MTT assay following the protocol previously described^33^. Cells were seeded into 96-well plates at a density of 3000 cells per well and treated with the indicated compounds for 72 h. MTT (MedChemExpress, HY-15924) reagent was then added to each well and incubated for 4 h at 37 °C to allow reduction of MTT to insoluble formazan crystals by metabolically active cells. Formazan crystals were subsequently solubilized, and absorbance was measured using a BioTek ELx808 microplate reader. Relative cell viability was calculated by normalizing absorbance values to vehicle-treated controls. Half-maximal inhibitory concentration (IC) values were determined by nonlinear regression analysis of dose-response curves using GraphPad Prism 9 (GraphPad Software).

### Quantification of fluorescence images

Identification and morphological analysis of ER-PM contact site puncta, as well as quantification of mRFP fluorescence intensity, were performed using custom MATLAB scripts. Fluorescence images were converted to grayscale and normalized to 16-bit intensity. Small punctate structures were enhanced using a morphological top-hat filter, followed by automated thresholding using Otsu’s method to generate binary masks. Morphological opening and size filtering were applied to remove noise and small artifacts. Puncta were further refined based on shape parameters, including circularity and eccentricity, to exclude elongated or irregular objects. The resulting masks were applied to the original images to extract puncta signals, and processed images were exported for downstream quantitative analysis.

Total contact site area fraction was calculated as the total area of detected contact site puncta divided by the total cell area for each cell. Average contact site size was defined as the mean area of individual contact site puncta per cell. Contact site density was calculated as the number of detected contact site puncta per cell divided by the total cell area. For withdrawal assays, the normalized total contact sites area fraction at each time point was calculated by dividing the contact sites total area fraction by the corresponding value at time 0. mRFP fluorescence intensity was quantified as the mean fluorescence intensity across the entire cell. Total contact site fluorescence intensity was calculated as the product of the mean mRFP fluorescence intensity and the total contact site area. Total cellular fluorescence intensity was calculated as the product of the mean mRFP fluorescence intensity and the total cell area.

To characterize the temporal kinetics of ER-PM contact sites formation, total contact site area fraction at each time point after ABA/Rapamycin addition or blue light illumination was fitted using single exponential association model, following the equation:

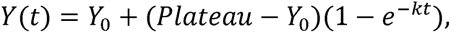

where *Y_0_*represents the initial contact-site area fraction at *t* = 0, Plateau denotes the maximal steady-state area reached at long time points, and *k* is the rate constant, which was used to derive the half-time of induction (*t*_1/2,on_ = ln2 / *k*).

For the recovery process, time-course data of normalized total contact site area fraction were fitted using single exponential dissociation model, following the equation:

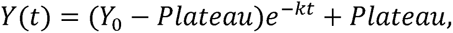

where *Y* represents the initial value at the start of recovery (t = 0), Plateau denotes the final steady-state value after complete dissociation, and *k* represents the rate constant used to calculate the half-time of recovery (*t*_1/2,off_ = ln2 / *k*).

### Statistical analysis

All data visualization and statistical analyses were performed using GraphPad Prism 9. Data shown in the figures are presented as mean ± s.e.m. For boxplots, the center line represents the median, boxes indicate the interquartile range (first to third quartiles), and whiskers denote the minimum and maximum values excluding outliers, and the plus sign (+) marks the mean. Comparisons between two groups were performed using two-tailed unpaired Student’s *t* tests unless otherwise specified. A *p* value less than 0.05 was considered statistically significant. In all figures, significance is coded as: ns non-significant, \**p* < 0.05, \*\**p* < 0.01, \*\*\**p* < 0.001, \*\*\*\**p* < 0.0001.

## RESULTS AND DISCUSSION

### An ABA-inducible CID system enables non-toxic, tunable, and reversible control of ER-PM contact site formation

To overcome the limitations of currently available rapamycin-based ER-PM contact-forming CID systems, including irreversibility, cytotoxicity, interference with endogenous mTOR signaling, and reduced specificity due to competition with endogenous FKBP or cyclophilin family proteins in eukaryotic cells, we sought to develop a novel abscisic acid (ABA)-inducible CID system to chemically control ER-PM contact site formation. CID systems based on ABA-induced heterodimerization between ABIcs- and PYLcs-tagged proteins represent a potentially advantageous alternative to rapamycin-based systems, as they are derived from plant signaling pathways and are therefore expected to exhibit reduced interference with host-cell signaling and improved molecular specificity^31^. Guided by the design of a previously established rapamycin-inducible ER-PM contact-forming system based on FRB/FKBP interactions, we implemented an analogous ER-PM tethering strategy using the ABIcs/PYLcs pair to construct an ABA-inducible ER-PM contact-forming CID system. A rapamycin-inducible ER-PM tethering system was constructed in parallel as a reference. In both systems, identical PM- and ER-targeting sequences were used to localize the dimerization domains to opposing membranes, such that ER-PM tethering differed only in the inducible dimerization module. For PM targeting, the N-terminal palmitoylation/myristoylation signal sequence of human Lyn kinase^34^ was fused to the N terminus of ABIcs or FRB tagged with mRFP. For ER targeting, the C-terminal ER localization sequence of human Sac1 phosphatase^22^ was fused to the C-terminus of PYLcs or FKBP tagged with mStaygold. In both systems, the PM- and ER-targeted fusion proteins were encoded within a single bicistronic expression cassette and separated by a self-cleaving T2A peptide^35^ (**Figure 1A**).

**Figure 1.**
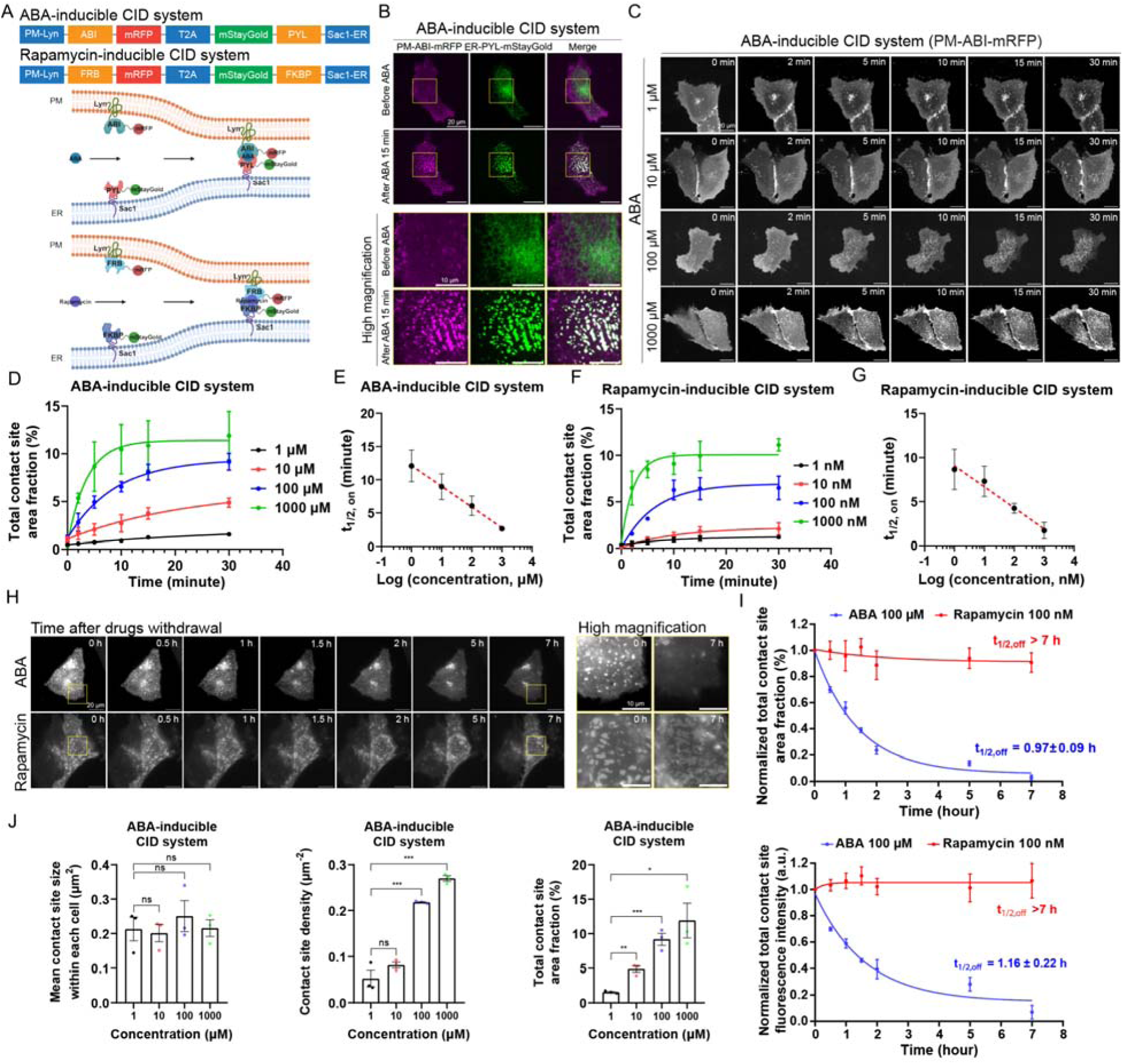
(**A**) Schematic representation of construct designs (top), the ABA-inducible CID system (middle), and the rapamycin-inducible CID system (bottom). (**B**) Representative confocal images of U2OS cells expressing the ABA-inducibleconstruct (PM-ABIcs-mRFP-T2A-mStayGold-PYLcs-ER) before and 15 min after treatment with 100 μM ABA. Yellow boxed regions are shown at higher magnification below. Scale bar: 20 μm (original view); 10 μm (magnified view). (**C**) Representative confocal images illustrating the time-course (0-30 min) and dose-dependent formation of ABA-induced ER-PM contact sites in U2OS cells. Scale bars: 20 μm. (**D**) Quantification of total ER-PM contact site area fraction from (**C**), fitted with single exponential association model. Data represent mean ± s.e.m. from three independent biological replicates (n = 3). (**E**) Half-time of ER-PM contact site formation (t_1/2,on_) calculated from the data shown in Fig. 1D, using single exponential association model. (**F**) Quantitative analysis of total ER-PM contact site area fraction from the time-course (0-30 min) and dose-response formation of rapamycin-induced ER-PM contact sites in U2OS cells (data from Fig. S3), fitted with single exponential association model. Data represent mean ± s.e.m. from three independent biological replicates (n = 3). (**G**) Half-time of ER-PM contact site formation (t_1/2,on_) calculated from the data shown in Fig. 1F, using single exponential association model. (**H**) Representative epi-fluorescence images showing the time-course (0-7 h) of U2OS cells following drug withdrawal after 30 min treatment with 100 μM ABA (top) or 100 nM rapamycin (bottom). Magnified views of the yellow boxed regions are shown on the right. Scale bars: 20 μm (original view); 10 μm (magnified view). (**I**) Quantification of the normalized total contact site area fraction (up) and normalized total contact sites fluorescence intensity (bottom) from Fig. 1H, fitted with single exponential dissociation model to calculate t_1/2,off_. Data represent mean ± s.e.m. from three independent biological replicates (*n* = 3). (**J**) Quantitative analysis of the mean size, density, and total area fraction of ER-PM contact sites in U2OS cells 30 min after treatment with increasing concentrations of ABA (1, 10, 100, and 1000 μM). Data represent mean ± s.e.m. from three independent biological replicates (*n* = 3). P values were calculated using two-tailed unpaired Student’s t-test: *\*P < 0.05, **P < 0.01, ***P < 0.001, ****P < 0.0001*, and ns, not significant.

We first validated whether the dimerization domains were correctly localized to their intended subcellular compartments and whether ABA could induce ER-PM contact site formation in the designed ABA-inducible ER-PM contact-forming CID system (PM-ABIcs-mRFP-T2A-mStayGold-PYLcs-ER), using confocal microscopy. Prior to ABA addition, U2OS cells expressing this construct showed uniform plasma membrane localization of PM-ABIcs-mRFP, while mStayGold-PYLcs-ER displayed an ER network pattern, with minimal colocalization between the two fluorescence signals (**Figure 1B**). Upon ABA addition, PM-ABIcs-mRFP and mStayGold-PYLcs-ER rapidly reorganized into discrete punctate structures at the cell surface within 5 min, where they exhibited tight signal co-localization (**Figure 1B**). These puncta appeared as bRight: well-defined foci with sharp boundaries, clearly distinct from the surrounding PM and ER regions. U2OS cells expressing the rapamycin-inducible ER-PM contact-forming CID system (PM-FBR-mRFP-T2A-mStayGold-FKBP-ER) exhibited similar localization patterns before stimulation and comparable puncta formation following rapamycin treatment (**Figure S1**), consistent with previous reports of rapamycin-induced ER-PM tethering^22^. Together, these results demonstrate that the ABA-inducible CID system enables robust and specific induction and visualization of ER-PM contact sites, with behavior comparable to that of the established rapamycin-inducible system^22^.

Next, we compared the cytotoxicity of ABA and rapamycin treatments, as ABA is not expected to interfere with endogenous mammalian signaling pathways, unlike rapamycin. U2OS cells were treated with increasing concentrations of ABA or rapamycin for 72 h, and cell viability was assessed using MTT assays. Rapamycin caused a dose-dependent decrease in viability (IC_50_ = 0.716 ± 0.358 μM), indicating substantial cytotoxicity, whereas ABA showed no detectable cytotoxic effects even at 1000 μM, with viability comparable to untreated controls (**Figure S2A-C**). The absence of cytotoxicity in the ABA system is consistent with its plant origin: ABA is a plant-derived phytohormone and ABA signaling is absent in mammalian cells, eliminating endogenous receptors or competing proteins. By contrast, rapamycin is a macrolide compound that potently inhibits the mTOR signaling pathway and exhibits well-documented cytostatic and cytotoxic effects across multiple mammalian cell types. Previous studies have shown that rapamycin induces growth inhibition or apoptosis in cell lines, such as HeLa, HEK293, U2OS, and MIN6, that are commonly used in ER-PM contact site studies^28,29,36,37^. Furthermore, because the mTOR-rapamycin-FKBP12 pathway is endogenous to mammalian cells, competition from endogenous FKBP12 may reduce the efficiency of exogenous FKBP-fused constructs, potentially compromising dimerization specificity and experimental reproducibility.

To quantify the kinetics of ER-PM contact site formation and assess whether it can be precisely controlled by ABA concentration, we monitored the time course of contact site formation at multiple ABA doses time (**Figure 1C**). During induction, the PM-anchored ABIcs-mRFP signal transitioned from a uniform plasma membrane distribution to discrete puncta, providing higher contrast than the mStayGold-PYLcs-ER signal, which reorganized from a reticular ER network and could lead to misidentification of ER structures as puncta. Therefore, PM-ABIcs-mRFP was used as the representative imaging channel for quantifying ER-PM contact sites. At each ABA concentration, the total area fraction of ER-PM contact sites, defined as the total contact site area per cell normalized to total cell area, increased exponentially over time (**Figure 1D**). The induction half-time (*t*_1/2,on_) was extracted by fitting the data to a single exponential function. Kinetic analysis revealed a clear dose dependence of *t*_1/2,on_. Specifically, as ABA concentration increased logarithmically (1, 10, 100, and 1000 μM), *t*_1/2,on_ decreased linearly (**Figure 1E**), indicating an ABA concentration-dependent acceleration of ER-PM contact site formation. A similar concentration-dependent kinetic behavior was observed for the rapamycin-induced CID system, in which *t*_1/2,on_ decreased linearly with increasing rapamycin concentrations (1-1000 nM) (**Figure 1F, G, Figure S3**). Importantly, this linear relationship between inducer concentration and induction kinetics enables precise and tunable control of ER-PM contact site formation: using the predetermined calibration curves (**Figure 1E,G**), a defined chemical concentration can be applied to achieve a desired rate of contact site formation.

This inducer (*i.e.*, ABA, rapamycin) concentration-dependent acceleration may arise from mass-action kinetics^38,39^, whereby increasing small-molecule inducer concentrations enhance the effective association rate of the CID pairs, or from receptor occupancy effects that yield a log-linear response to inducer concentration^40^. Additionally, the trend may reflect modulation of a rate-limiting nucleation step during ER-PM contact site assembly. The concurrent decrease in *t*_1/2,on_ and increase in steady-state total contact site area fraction at higher inducer concentrations suggest that ABA or rapamycin primarily regulates the frequency and probability of nucleation events rather than the lateral expansion rate of individual contact zones. At elevated inducer concentrations, rapid formation of ABIcs/PYLcs or FRB/FKBP complexes likely increases the density of productive nucleation events at the ER-PM interface, leading to faster and more extensive contact site assembly. Together, these findings demonstrate that both the extent and kinetics of chemically induced ER-PM contact site formation scale linearly with the inducer concentration. Importantly, this tunable and predictable dose-response relationship enables precise quantitative control of ER-PM contact dynamics, establishing a calibration-like framework analogous to analytical standard curves for dissecting chemically induced inter-organelle tethering dynamics.

To evaluate the reversibility of chemically induced ER-PM contact site formation, we examined and compared the recovery kinetics of the ABA- and rapamycin-based CID systems following inducer withdrawal. U2OS cells expressing either system were first treated with the corresponding inducer to fully establish ER-PM contact sites, after which the inducer was removed by washing the cells five times with fresh inducer-free medium. Time-lapse imaging showed that ABA-induced ER-PM contact sites gradually disassembled and returned to baseline levels within ∼7 h after ABA removal (**Figure 1H**). In contrast, identical washout conditions failed to reverse rapamycin-induced ER-PM contacts over the same time period, indicating a marked difference in reversibility between the two systems. Quantification of the total ER-PM contact site area fraction and total contact site fluorescence intensity per cell revealed recovery half-times (*t*_1/2,off_) of 0.97 ± 0.09 h for total contact-site area fraction and 1.16 ± 0.22 h for total contact site fluorescence intensity (**Figure 1I**) in the ABA system, whereas rapamycin-induced contacts showed *t*_1/2,off_ values exceeding 7 h. These differences likely reflect fundamental differences in molecular affinity and complex stability of the CID pairs. The ABIcs/ABA/PYLcs interaction exhibit lower affinity and faster dissociation kinetics^31^ than the high-affinity FRB/rapamycin/FKBP ternary complex^26,41,42^, rendering ABA-induced ER-PM contacts readily reversible. This reversibility provides significant experimental advantages, including the ability to perform repeated induction-recovery cycles, dissect the kinetics of contact site nucleation versus expansion, and transiently perturb ER-PM contact-dependent processes without inducing long-lasting structural alterations.

Having established that the ABA-inducible system exhibits superior reversibility and nontoxicity compared to the rapamycin-inducible system, we next examined how steady-state ER–PM contact-site geometry responds to ABA concentration. Cells were treated with increasing ABA doses (1, 10, 100, and 1000 μM), and contact sites were allowed to reach steady state after 30 min of induction. We then quantitatively analyzed three geometric parameters: the mean contact-site size (average area of individual contacts within each cell), contact-site density (number of contacts per unit cell area), and the total contact-site area fraction per cell. As ABA concentration increased logarithmically, both contact-site density and total contact-site area fraction increased monotonically, whereas the mean contact-site size remained largely unchanged across the tested concentration range (**Figure 1J**). These results suggest that ABA concentration primarily controls ER-PM contact-site nucleation and abundance, whereas the lateral dimensions of individual contacts are constrained by the underlying tether architecture rather than inducer dose. Together, our results demonstrate that the ABA-inducible CID system developed here, in contrast to rapamycin-based CID systems, enables nontoxic, tunable, and reversible manipulation and visualization of ER-PM contact sites with minimal interference from endogenous cellular pathways. These properties make the ABA system particularly well suited for dynamic live-cell imaging, quantitative biophysical analysis, and reversible control of inter-organelle interactions.

### An optogenetic LID system enables rapid, tunable, and reversible control of ER-PM contact site formation

Though the ABA-inducible CID system developed here provides nontoxic, tunable, and reversible control of ER-PM contact site formation and visualization, the recovery kinetics remain relatively slow with a recovery half-time (*t*_1/2,off_) of ∼1 hour. To further accelerate reversibility, we turned to optogenetic tools, which enable genetically encoded, light-responsive modulation of protein interactions with high temporal precision in living cells. We therefore designed a dual-component light-induced dimerization (LID) system based on iLID/SspB pair to induce ER-PM contact site formation, where the iLID module is derived from the LOV2 domain of *Avena sativa* phototropin 1 and has been engineered for robust, rapid, and reversible blue-light-induced heterodimerization with SspB^43,44^. Our design of light-inducible ER-PM contact site forming LID system employed the same PM and ER targeting strategy as the ABA-inducible CID system: iLID was fused to a PM-targeting peptide (N-terminal Lyn motif) and tagged with mRFP, and SspB was fused to an ER-localization peptide (C-terminal Sac1 sequence) and tagged with mStayGold. The two fusion proteins were expressed from a single plasmid as separate polypeptides using a self-cleaving T2A peptide. In the dark, the Jα helix of iLID remains docked against the LOV2 core, sterically masking the embedded SsrA peptide and preventing its association with SspB. Upon blue-light illumination, LOV2 undergoes a conformational change that triggers undocking of the Jα helix, exposing SsrA and allowing its binding to ER-localized SspB, which in turn drives the assembly of ER-PM contact sites (**Figure 2A**).

**Figure 2.**
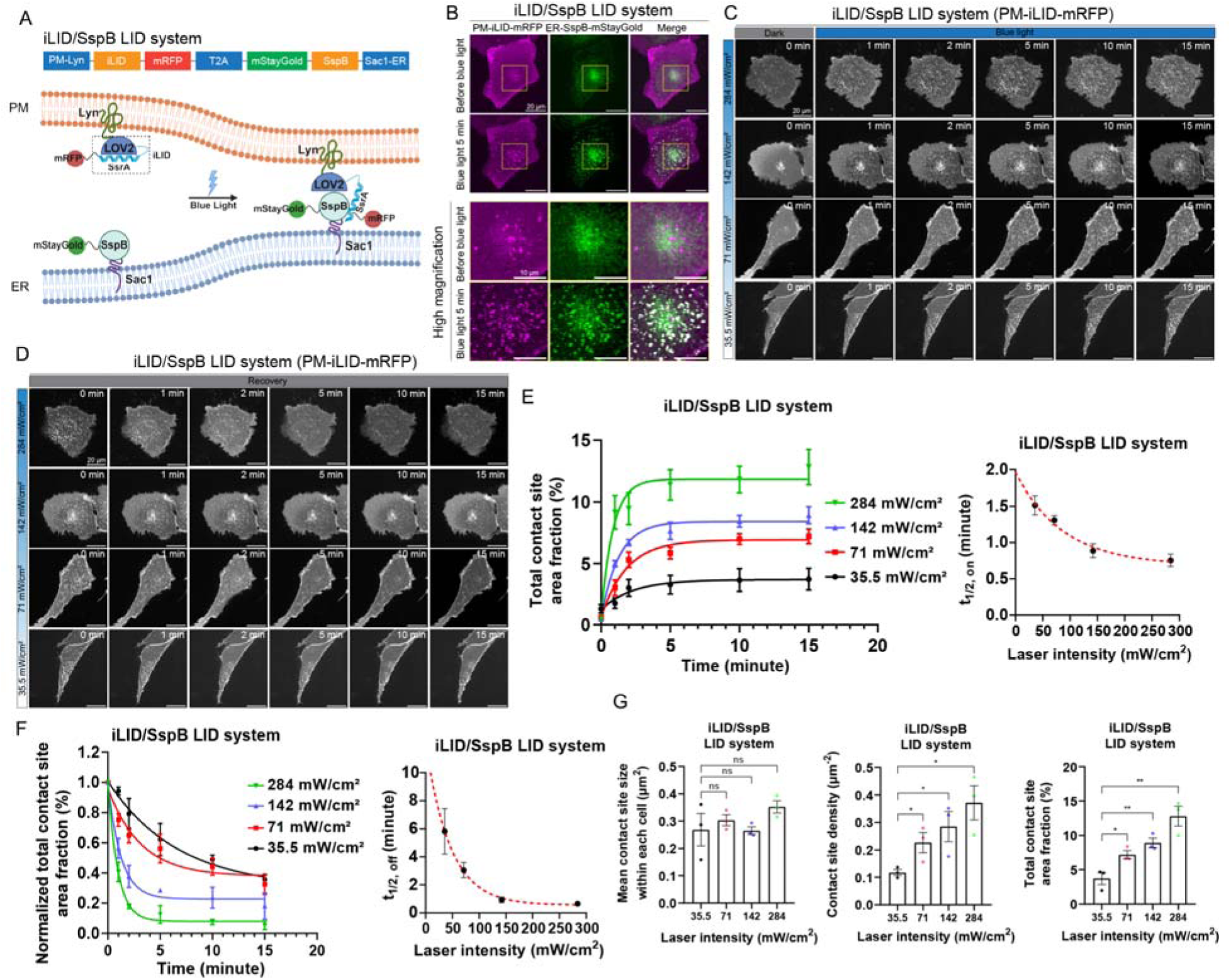
(**A**) Schematic representation of construct design (top) and the iLID/SspB LID system (bottom). (**B**) Representative confocal images of U2OS cells expressing the iLID/SspB LID construct (PM-iLID-mRFP-T2A-mStayGold-SspB-ER) before and after blue light stimulation (∼142 mW/cm^2^). Yellow boxed regions are shown at higher magnification below. Scale bar: 20 μm (original view); 10 μm (magnified view). (**C**, **D**) Representative confocal images showing the time-course (0-15 min) and light-intensity response of blue light-induced ER-PM contact site formation (**C**) and the subsequent recovery after light withdrawal (**D**) in U2OS cells. Blue light stimulation was applied using a 470-nm laser at 5-40% output, corresponding to 0.022-0.177 mW measured at the sample plane (power meter). Scale bar: 20 μm. (**E**) Quantification of total ER-PM contact site area fraction from Fig. 2C. Left: time-course of blue light-induced ER-PM contact sites formation. Data were fitted with single exponential association model. Right: half-time of ER-PM contact site formation (t_1/2,on_) calculated from the data shown in left using single exponential association model. Data represent mean ± s.e.m. from three independent biological replicates (*n* = 3). (**F**) Quantification of total ER-PM contact site area fraction from Fig. 2D. Left: recovery of ER-PM contact sites after blue light withdrawal. Data were fitted with single exponential dissociation model. Right: half-time of ER-PM contact site dissociation (t_1/2,off_) calculated from the data shown in left using single exponential dissociation model. Data represent mean ± s.e.m. from three independent biological replicates (*n* = 3). (**G**) Quantitative analysis of the mean size, density, and total area fraction of ER-PM contact sites in U2OS cells 15 min after treatment with increasing intensity of blue light stimulation. Data represent mean ± s.e.m. from three independent biological replicates (*n* = 3). P values were calculated using two-tailed unpaired Student’s t-test: *\*P < 0.05, **P < 0.01, ***P < 0.001, ****P < 0.0001*, and ns, not significant.

To validate whether the iLID/SspB system can effectively induce ER-PM contact site formation, we transfected U2OS cells with the construct PM-iLID-mRFP-T2A-mStayGold-SspB-ER and monitored contact site formation using confocal microscopy before and after light stimulation. In the dark, PM-iLID-mRFP fluorescence was uniformly distributed along the PM, whereas the SspB-mStayGold-ER exhibited an ER network pattern, with minimal background puncta formation, closely resembling the localization observed for the ABA-inducible CID system. Upon blue-light illumination (470 nm at 20% laser output, corresponding to an intensity of ∼142 mW/cm^2^ at the sample plane), both PM-iLID-mRFP and SspB-mStayGold-ER rapidly reorganized into discrete punctate structures at the cell surface, where they displayed tight co-localization. These punctate ER-PM contact sites formed rapidly in a time-dependent manner, reaching a near-steady state within 5 min, after which the puncta remained largely stable with minimal lateral expansion (**Figure 2B**). These results demonstrate that our designed light-inducible LID system enables robust and rapid induction and visualization of ER-PM contact sites.

Inspired by our earlier findings that ER-PM contact site formation can be precisely tuned by ABA or rapamycin concentration, and that ABA-induced contacts are reversible, we next tested whether light intensity could similarly control the formation and dissociation kinetics of our light-inducible ER-PM contact system. Increasing 470 nm laser intensity from 5% (∼35.5 mW/cm^2^) to 40% (∼284 mW/cm^2^) progressively accelerated contact-site formation and increased the steady-state contact-site area fraction (**Figure 2C**). Following 15 min of illumination at each tested laser intensity, withdrawal of blue light triggered rapid dissociation of ER-PM contact sites across all conditions, demonstrating robust reversibility (**Figure 2D**). Notably, higher illumination intensities resulted in faster dissociation upon light withdrawal. To quantify these effects, the time-dependent changes in total contact site area fraction during light-induced formation and dark-state dissociation were fitted with single-phase exponential association and dissociation functions, respectively (**Figure 2E,F**), yielding the induction and recovery half-times (*t*_1/2,on_ and *t*_1/2,off_). Both *t*_1/2,on_ and *t*_1/2,off_ decreased exponentially with increasing 470-nm laser intensity (**Figure 2E,F**), indicating that light intensity provides precise, bidirectional control over ER-PM contact-site kinetics. The coupled acceleration of both formation and dissociation at higher light intensities suggests a shared kinetic origin. Increased illumination rapidly drives a larger fraction of iLID molecules into the binding-competent conformation, enhancing the effective association rate and promoting rapid contact-site nucleation. Upon light withdrawal, these activated iLID molecules undergo photophysical relaxation to the dark, non-binding state, leading to a rapid loss of productive interactions and accelerated contact-site disassembly. This behavior is consistent with mass-action effects and a nucleation-dominated assembly mechanism, in which light intensity modulates the population dynamics of the activated iLID state rather than altering intrinsic binding affinity. Compared with the ABA-inducible CID system, the iLID/SspB-based LID system exhibited markedly faster dynamics, with rapid induction (5% laser output, *t*_1/2,on_ = 1.51 ± 0.13 min) and near-complete recovery within minutes (5% laser output, *t*_1/2,off_ = 5.82 ± 1.62 min). In contrast, the ABA system required ∼1 h for contact-site dissociation. Together, these results demonstrate the superior temporal reversibility of the iLID/SspB-based ER-PM contact site forming system, enabling repeated cycles of contact formation and dissociation with precisely tunable kinetics controlled by light intensity.

Next, we examined how steady-state ER-PM contact-site geometry responds to optical input in the iLID/SspB system. Cells were subjected to increasing 470 nm laser intensities, and ER-PM contacts were allowed to reach steady state after 15 min of continuous illumination. As in the ABA-inducible system, we quantified three geometric parameters: the mean size of individual contact sites, contact-site density, and the total contact-site area fraction per cell. Increasing bule laser intensity led to a monotonic increase in both contact-site density and total contact-site area fraction, whereas the mean contact-site size remained largely unchanged across the tested intensity range (**Figure 2G**). This behavior mirrors the inducer dose dependence observed for the ABA system, indicating that optical input primarily regulates ER-PM contact-site nucleation and overall abundance. In contrast, the lateral dimensions of individual contact sites appear to be constrained by the intrinsic tether architecture rather than the strength of the inducing stimulus.

It is worth noting that prior to this work, a light-inducible ER-PM contact site-forming system, termed LiMETER, had been reported^21,45^. However, our iLID/SspB-based LID system offers two key advantages. First, LiMETER consists of a single GFP-tagged component localized to the ER, in which blue-light activation of the LOV2 module exposes a polybasic region that binds PI(4,5)P_2_ at the PM to induce ER-PM contacts^21^. As a result, the LiMETER fluorescence signal largely follows the tubular and sheet-like ER network, making it difficult to visually distinguish discrete ER-PM contact-site puncta from the surrounding ER structure. In our hands, this led to frequent misidentification of ER network features as contact-site puncta by MATLAB-based image analysis, even in the absence of light stimulation (**Figure S4A,B**), thereby reducing contrast between dark and illuminated states (**Figure S4C**). Second, LiMETER relies on light-induced exposure of a polybasic motif that directly binds PI(4,5)P_2_, rendering ER-PM contact formation dependent on the local phosphoinositide environment and potentially perturbing endogenous PI(4)P/PI(4,5)P_2_ dynamics at contact sites. Such lipid-dependent membrane association may complicate studies aimed at dissecting the interplay between ER-PM contacts and lipid signaling. By contrast, our iLID/SspB system operates through lipid-independent protein-protein interactions, minimizing interference with native phosphoinositide signaling. This design enables robust, tunable, reversible ER-PM contact formation while preserving the physiological lipid landscape, providing a versatile optogenetic platform for precise spatiotemporal manipulation and visualization of ER-PM contacts.

### Systematic linker-length variation enables control of ER-PM contact-site stability and geometry

Emerging evidence suggests that not only the presence, but also the density, size, and nanoscale organization of ER-PM contact sites critically modulate local signaling and global cellular behavior^15–17^. For example, in migrating cells, ER-PM contacts display a pronounced front-back gradient in both density and size, with small, highly dynamic contacts at the leading edge and larger, more stable contacts at the trailing edge^15^. These observations highlight the functional importance of ER-PM contact site geometry and dynamics. Motivated by this, we sought to leverage the chemically induced dimerization (CID) and light-induced dimerization (LID) systems developed here to introduce precise control over the average size of induced ER-PM contact sites, as well as the integrated total ER-PM contact area per cell. Such tunability provides a powerful platform for dissecting geometry-dependent functions of ER-PM contacts across diverse cellular contexts, including polarized migration, signal activation, and mechanical remodeling, and offers a generalizable strategy for probing structure-function relationships at membrane contact sites with high spatial and temporal precision.

To achieve tunable control over ER-PM contact site geometry, including average size and total contact area per cell, we first tested whether varying the ER-PM intermembrane spacing at induced contact sites could modulate these geometry parameters. Specifically, we engineered constructs with variable intermembrane spacer lengths by modifying both the ABA-inducible and iLID/SspB-based light-inducible systems described above (**Figures 3A and 4A**). In these designs, rigid α-helical spacers composed of repeated (EAAAR)_n_ motifs^46^ (n=0, 4, 9) were inserted between the membrane-targeting sequences and the dimerization modules (*i.e.*, between the ER- or PM-localization sequence and the ABI or PYL component in the ABA-based system, and between the localization sequence and the iLID or SspB component in the light-inducible system). (EAAAR)_n_ peptides serve as model helices whose helicity is markedly superior to many rationally designed α-helical peptides^47,48^. Owing to their compact, stable, and highly directional α-helical structure, these peptides have been successfully applied in high-throughput screening of rapamycin-mediated FKBP12-FRB interactions^49^. Based on previously established length estimates for (EAAAR) α-helical spacers, in which each (EAAAR) module spans approximately 3 nm in length^45,46^, and previously established length estimates for the dimerization modules^22,45^, we estimated that our three designed constructs (referred to as (EAAAR)_0_, (EAAAR)_4_ and (EAAAR)_9_ hereafter) produce ER-PM intermembrane spacings of approximately ∼8-12 nm, ∼14-18 nm, and ∼20-24 nm, respectively. Together, these spacings effectively cover the full range of distances (∼10-30 nm) reported for native ER-PM contact sites^50^.

**Figure 3.**
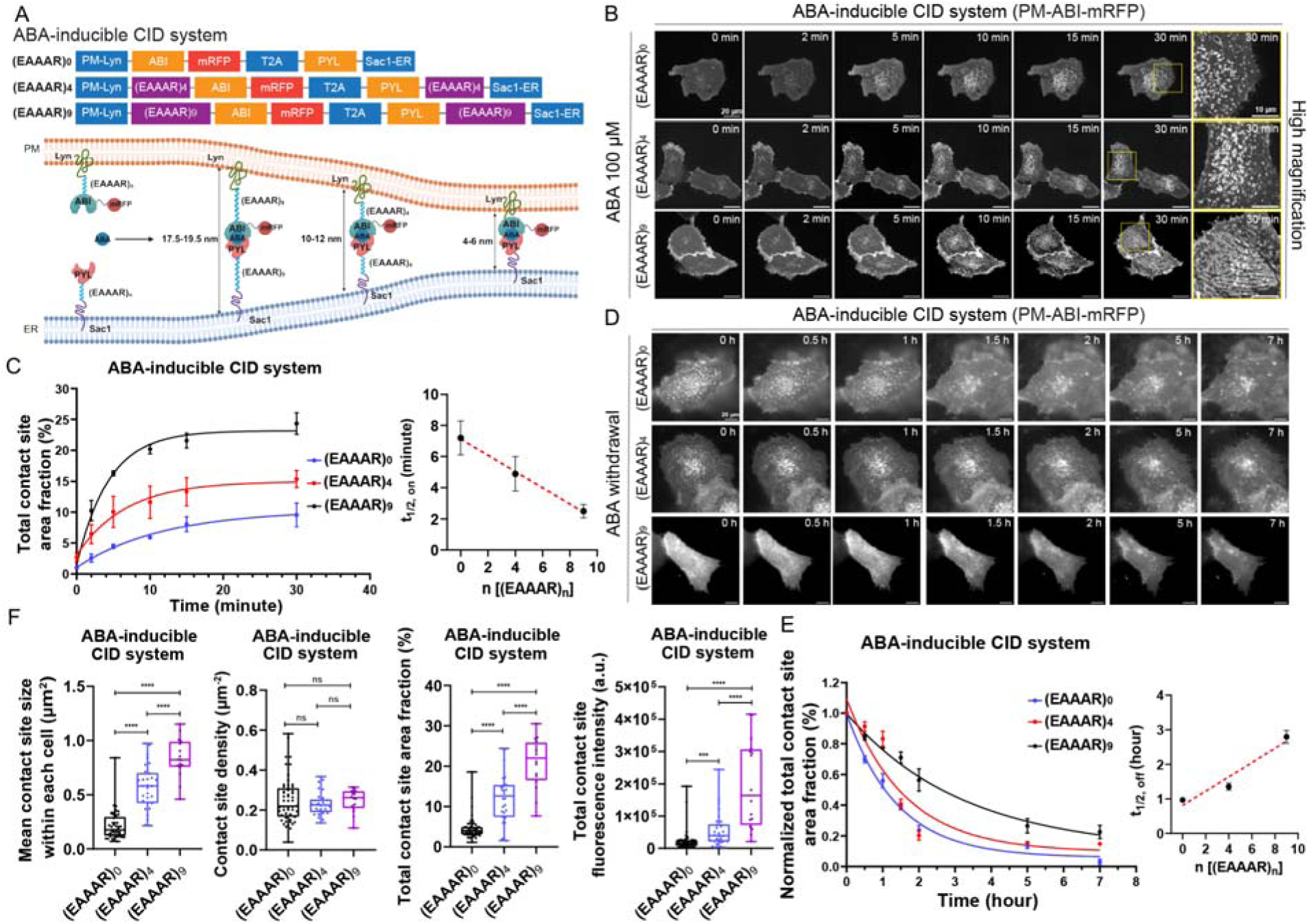
(**A**) Schematic showing construct designs (top) and the use of the ABA inducible CID system with varying spacer lengths to manipulate inter-membrane distances at ER-PM contact sites (bottom). (**B**) Representative time-course (0-30 min) and spacer-response images of 100 μM ABA-induced ER-PM contact site formation in U2OS cells. Cells were transfected with different spacer constructs and imaged by confocal microscopy 48 h after transfection. Yellow boxed regions in the 30-min group are shown at higher magnification on the right. Scale bars: 20 μm (original view); 10 μm (magnified view). (**C**) Quantification of total ER-PM contact site area fraction derived from the images shown in Fig. 3B. Left: time-course data fitted with single exponential association model. Right: half-time of ER-PM contact site formation (t_1/2,on_) calculated from the time-course data using single exponential association model in GraphPad Prism. Data represent mean ± s.e.m. from three independent biological replicates (*n* = 3). (**D**) Representative time-course imaging (0-7 h) of U2OS cells following ABA withdrawal after 30 min of treatment, across different spacer groups. Images were acquired using epi-fluorescence microscopy. Scale bars: 20 μm. (**E**) Quantification of normalized total contact site area fraction from Fig 3D. Left: time-course data fitted with single exponential dissociation model. Right: half-time of ER-PM contact site dissociation (t_1/2,off_) calculated from the time-course data using single exponential dissociation model. Data represent mean ± s.e.m. from three independent biological replicates (*n* = 3). (**F**) Quantitative analysis of the mean size, density, total area fraction, and total fluorescence intensity of ER-PM contact sites in U2OS cells 15 min after 100 μM ABA treatment across different spacer constructs: (EAAAR)_0_ (*n* = 51 cells), (EAAAR)_4_ (*n* = 30 cells), and (EAAAR)_9_ (*n* = 16), pooled from three independent biological replicates. P values were calculated using two-tailed unpaired Student’s t-test: *\*P < 0.05, **P < 0.01, ***P < 0.001, ****P < 0.0001*, and ns,not significant.

**Figure 4.**
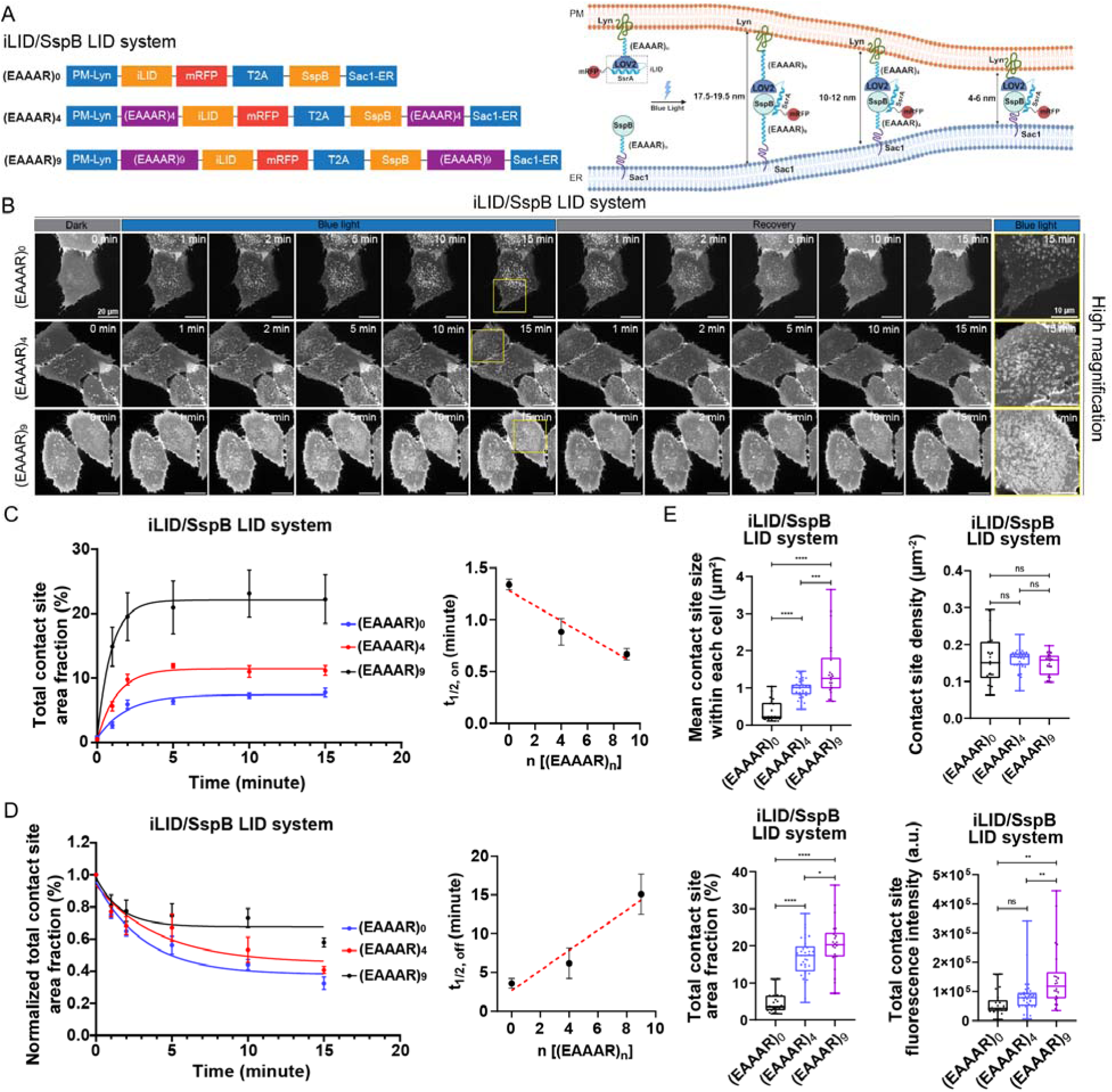
**(A)** Schematic showing construct designs (left) and the use of the iLID/SspB LID system with varying spacer lengths to manipulate inter-membrane distances at ER-PM contact sites (right). **(B)** Representative time-course images of ER-PM contact site dynamics in U2OS cells transfected with different spacer constructs. Cells were imaged by confocal microscopy 48 h after transfection during blue light-induced contact site formation (0-15 min, on; 71 mW/cm^2^) and following blue light withdrawal (0-15 min, off) after 15 min of illumination. Yellow boxed regions in the 15-min group (on) are shown at higher magnification on the right. Scale bar: 20 μm (original view); 10 μm (magnified view). **(C)** Left: quantification of total ER-PM contact site area fraction from Fig. 4B during blue light-induced contact site formation (on), fitted with single exponential association model. Right: half-time of ER-PM contact site formation (t_1/2,on_) calculated from left using single exponential association model. Data represent mean ± s.e.m. from three independent biological replicates (*n* = 3). **(D)** Left: quantification of normalized total ER-PM contact site area fraction from Fig. 4B following blue light withdrawal after 15 min illumination (off), fitted with single exponential dissociation model. Right: half-time of ER-PM contact site dissociation (t_1/2,off_) calculated from left using single exponential dissociation model. Data represent mean ± s.e.m. from three independent biological replicates (*n* = 3). **(E)** Quantitative analysis of mean size, density, total area fraction, and total fluorescence intensity of ER-PM contact sites in U2OS cells after 5 min blue light illumination (71 mW/cm^2^) across different spacer constructs: (EAAAR)_0_ (*n* = 22 cells), (EAAAR)_4_ (*n* = 35 cells), and (EAAAR)_9_ (*n* = 22 cells), pooled from three independent biological replicates. P values were calculated using two-tailed unpaired Student’s t-tests: *\*P < 0.05, **P < 0.01, ***P < 0.001, ****P < 0.0001*, and ns, not significant.

We first examined whether the engineered (EAAAR)_0_, (EAAAR)_4_ and (EAAAR)_9_ constructs, which differ in linker (*i.e.*, spacer) length, exhibit distinct kinetics of ER-PM contact formation and disassembly. U2OS cells expressing these constructs were subjected to time-course imaging following induction with 100 μM ABA (for the ABA-inducible CID system) or 10% blue-light illumination (for the light-inducible iLID/SspB system; 10% laser output corresponding to ∼71 mW/cm^2^ at the sample plane). In both systems, all three constructs exhibited robust, time-dependent ER-PM contact site formation (**Figure 3B and 4B**). We quantified the total ER-PM contact site area fraction per cell as a function of time after stimulation. Fitting the induction time traces with single exponential association model yielded half-times of contact formation (t_1/2,on_), which decreased linearly with increasing linker length (**Figure 3C and 4C**). We next assessed the disassembly kinetics of the three linker variants by washing out ABA (for the ABA-inducible CID system) or terminating blue-light illumination (for the light-inducible iLID/SspB system) (**Figure 3D and 4B**). Recovery time traces were fit using a single-phase exponential dissociation model to extract the half-times of recovery (t_1/2,off_). In contrast to the formation kinetics, t_1/2,off_ increased with linker length in both systems (**Figure 3E and 4D**). Notably, the constructs containing the longer linker exhibited a substantially elevated residual plateau, corresponding to incomplete recovery within the observation time window. This behavior suggests the emergence of a more persistent contact population that resists rapid disassembly. Together, these results indicate that linker length is a key determinant of both ER-PM contact site formation and disassembly kinetics. The reduced recovery observed for longer linkers likely reflects a transition from predominantly monovalent, kinetically labile tethering to a multivalent, avidity-stabilized state. Extended spacers permit larger contact footprints and enable simultaneous tethering interactions, thereby increasing local binding density, promoting cooperative recruitment, and raising the energetic barrier for dissociation. In addition, longer linkers may facilitate local membrane flattening and lateral clustering of tethering complexes, further stabilizing contacts and slowing recovery through increased viscous dissipation.

It is important to note that in contrast to tuning ABA concentration or iLID/SspB light power, which accelerates both contact formation (decreasing t_1/2,on_) and disassembly (decreasing t_1/2,off_) (**Figure 1E** and **2E,F**), linker-length modulation accelerates formation but stabilizes ER-PM contacts, resulting in increased t_1/2,off_. This difference arises because ABA or light intensity controls the fraction of active tethers, enhancing nucleation and turnover without changing intrinsic tether avidity, whereas linker length alters the physical reach and multivalency of tether interactions, directly influencing contact stability. This linker-length-dependent modulation provides a unique means to tune ER-PM contact stability, rather than merely controlling formation and disassembly dynamics.

Next, we investigated whether varying linker length could systematically tune ER-PM contact site geometry, as predicted. U2OS cells expressing (EAAAR)_0_, (EAAAR)_4_, and (EAAAR)_9_ constructs were stimulated with 100 μM ABA for 15 min (ABA-inducible CID system) or 10% (∼71 mW/cm^2^) blue-light illumination for 5 min (for the light-inducible iLID/SspB system), followed by cell fixation, confocal imaging and quantitative analysis. In both systems, the mean size, total area fraction, and total fluorescence intensity of induced ER-PM contact sites per cell increased monotonically with linker length, whereas contact site density remained largely unchanged (**Figure 3F and 4E**). Consistent with these quantitative trends, cells expressing the long-linker (EAAAR)_9_ variant exhibited large, island-like contact sites in contrast to the discrete small puncta observed for the (EAAAR)_0_ construct (**Figure 3B and 4B**). These results demonstrate that linker elongation provides a precise and robust means to tune the average size of individual ER-PM contact sites without substantially altering contact site density, thereby increasing the total ER-PM contact area per cell across both chemically and optically inducible tethering platforms. Notably, linker-length–dependent tuning selectively regulates contact-site stability and lateral growth, in contrast to inducer-dose–dependent regulation (ABA concentration or laser intensity), which predominantly controls contact-site nucleation and abundance rather than the lateral growth of individual contacts.

Collectively, these results establish linker length as a quantitative and predictive design parameter that couples molecular architecture to ER-PM contact-site stability and geometry. By systematically tuning linker length, we can independently modulate contact-site stability, steady-state size, and total contact area without substantially altering contact density. This reveals that the physical reach of inducible tethers directly governs the balance between kinetically labile and avidity-stabilized contact states. Such tunable behavior defines explicit structure-function relationships for membrane contact sites and enables rational engineering of contacts with predefined stabilities and geometries.

### Systematic tether-abundance variation enables orthogonal control of ER-PM contact-site geometry

In addition to varying the linker length (i.e., intermembrane distance), modulating the concentrations of the tethering components on the ER and PM membranes may provide an alternative and orthogonal means of tuning ER-PM contact-site geometry. This is because the local density of membrane-anchored tethers directly determines the probability of productive tether-tether encounters, the extent of lateral recruitment following nucleation, and the degree of cooperative stabilization within a contact zone. Consequently, altering tether abundance is expected to influence key geometric parameters of ER-PM contacts without changing the intrinsic intermembrane spacing imposed by tether (*i.e.*, linker) length.

To test this hypothesis, we selected the long-linker (EAAAR)_9_ construct and subcloned it into a previously established XLone backbone containing a doxycycline (Dox)-inducible promoter ^51^ for both the ABA-inducible CID system (hereafter referred to as XLone-ABA) and light-inducible iLID/SspB system (hereafter referred to as XLone-iLID/SspB) (**Figure 5A**). The XLone system combines a Tet-On 3G promoter and PiggyBac transposon elements, enabling stable genomic integration and tightly regulated, Dox-dependent expression^51^. This configuration minimizes transgene silencing and supports reversible, tunable control of gene expression, making it particularly suitable for precisely adjusting the expression levels of tethering components on the ER and PM membranes.

**Figure 5.**
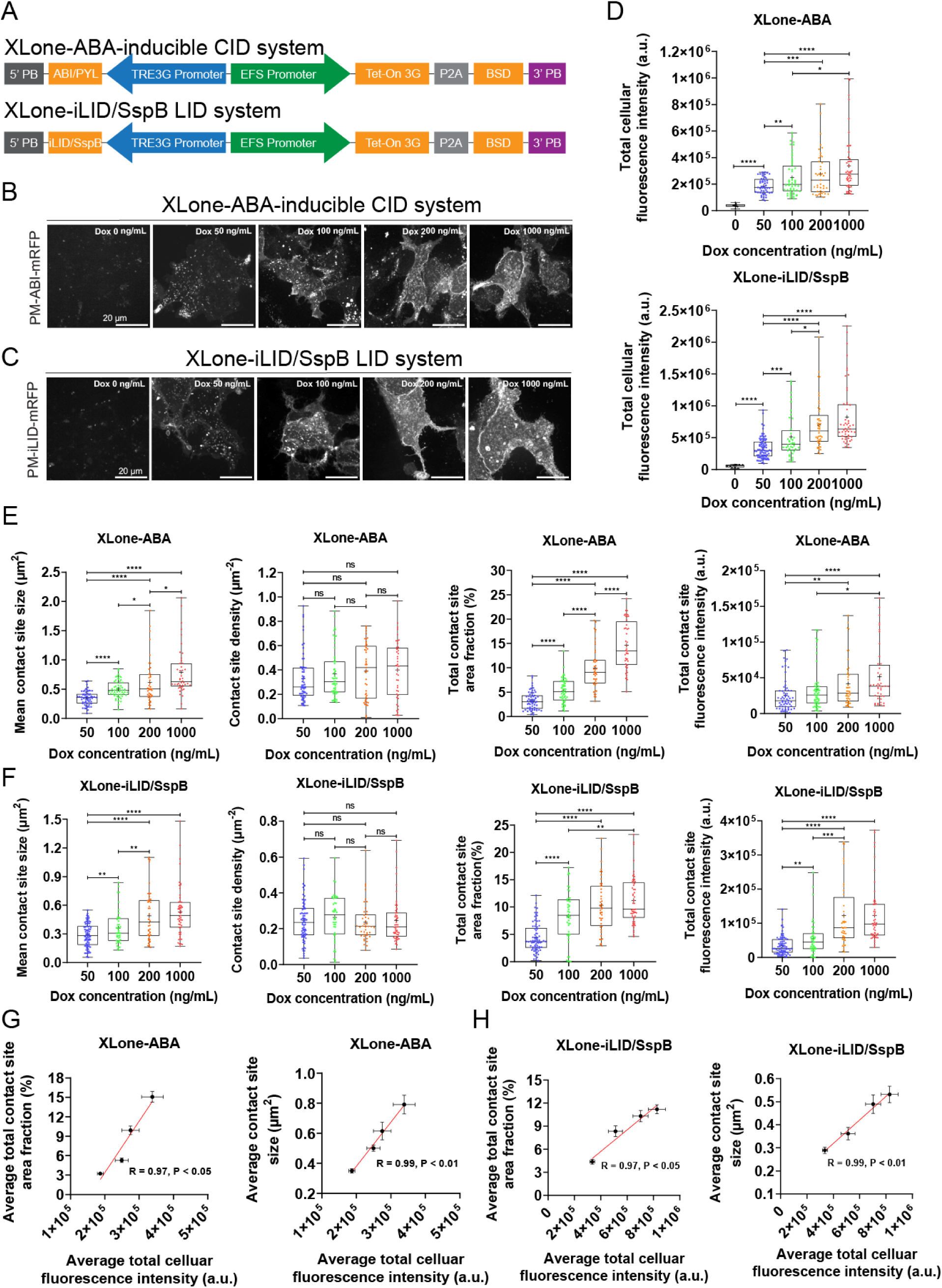
(**A**) Schematic showing XLone-ABA-inducible CID system and XLone-iLID/SspB LID system construct designs. 5′ PB, 5′ PiggyBac Terminal Repeat; 3’ PB, 3’ PiggyBac Terminal Repeat; BSD, Blasticidin resistance gene; P2A, porcine teschovirus-1 2A self-cleaving peptide. (**B, C**) Representative confocal images of ER-PM contact sites in HEK293T cells transfected with XLone-ABA-inducible CID construct (**B**) or XLone-iLID/SspB LID construct (**C**), pre-induced with increasing doxycycline concentrations (0-1000 ng/mL). Cells were subsequently treated with 100 μM ABA for 30 min (**B**) or illuminated with blue light (4 mW/cm^2^) for 15 min (**C**). Scale bars: 20 μm. (**D**) Quantitative analysis of total cellular fluorescence intensity in HEK293T cells pre-induced with doxycycline at the indicated concentrations and subsequently stimulated with ABA (top, 100 μM, 30 min) or blue light (bottom, 15 min). For ABA-inducible CID system: 0 ng/mL (*n* = 8), 50 ng/mL (*n* = 49), 100 ng/mL (*n* = 52), 200 ng/mL (*n* = 43), 1000 ng/mL (*n* = 48); for iLID/SspB LID system: 0 ng/mL (*n* = 10), 50 ng/mL (*n* = 81), 100 ng/mL (*n* = 41), 200 ng/mL (*n* = 46), 1000 ng/mL (*n* = 53). Data were pooled from three independent biological replicates. **(E)** Quantitative analysis of the total area fraction, average size, density, and total fluorescence intensity of ER-PM contact sites in HEK293T cells pre-induced with doxycycline at the indicated concentrations, followed by treatment with 100 μM ABA for 30 min. Doxycycline concentrations were 50 ng/mL (*n* = 77 cells), 100 ng/mL (*n* = 44 cells), 200 ng/mL (*n* = 44 cells), and 1000 ng/mL (*n* = 55 cells), pooled from three independent biological replicates. **(F)** Quantitative analysis of the mean size, density, total area fraction, and total fluorescence intensity of ER-PM contact sites in HEK293T cells pre-induced with doxycycline at the indicated concentrations, followed by treatment with blue light (4 mW/cm^2^) for 15 min. Doxycycline concentrations were 50 ng/mL (*n* = 61 cells), 100 ng/mL (*n* = 55 cells), 200 ng/mL (*n* = 43 cells), and 1000 ng/mL (*n* = 44 cells), pooled from three independent biological replicates.(**G**, **H**) Correlations between average total cellular fluorescence intensity and ER-PM contact site parameters, including average total contact site area fraction (left) and average contact site size (right), in ABA-inducible CID (**G**) or iLID/SspB LID (**H**) systems. Pearson correlation coefficients (R) and corresponding P values were calculated using GraphPad Prism 9. P values were calculated using two-tailed unpaired Student’s t-tests: *\*P < 0.05, **P < 0.01, ***P < 0.001, ****P < 0.0001*, and ns, not significant.

We transfected HEK293T cells with the XLone-ABA and XLone-iLID/SspB constructs. 24 hours after transfection, cells were treated with a graded series of Dox concentrations (50, 100, 200, and 1000 ng/mL) to induce distinct expression levels of the tethering components on the ER and PM membranes. After an additional 24 hours of Dox induction, cells were stimulated with 100 μM ABA for 30 min (ABA-inducible CID system) or ∼4 mW/cm^2^ blue-light illumination for 15 min (for the light-inducible iLID/SspB system), followed by cell fixation, confocal imaging and quantitative analysis. In both systems, mRFP fluorescence intensity, which reports the expression level of the PM-localized tether (PM-ABIcs-mRFP for ABA-inducible CID system or PM-iLID-mRFP for light-inducible iLID/SspB system), increased monotonically with Dox concentration (**Figure 5B-D**). Concomitantly, multiple geometry-related parameters of induced ER-PM contact sites, including the mean contact-site size, total contact-site area fraction, and total contact-site fluorescence intensity per cell, increased in a Dox-dependent manner, whereas contact-site density remained largely unchanged (**Figure 5E, F**), consistent with the linker-length–dependent tuning we observed. Population-averaged analysis further revealed linear relationships between (i) average contact-site size and average PM-tether expression level, (ii) average contact-site density and average PM-tether expression level, (iii) average total contact-site area fraction and average PM-tether expression level, and (iv) average total contact-site fluorescence intensity and average PM-tether expression level (average PM-tether expression was quantified by total cellular mRFP fluorescence; **Figure 5G, H, and Figure S5A, B**). These results demonstrate that tuning the expression level of membrane-anchored tethering components via Dox induction provides a robust and quantitative means to modulate ER-PM contact-site geometry, like tuning the linker length.

Together, these results establish tether concentration as an independent and orthogonal design parameter for controlling ER-PM contact-site geometry. Whereas linker (tether) length primarily sets the intermembrane spacing and influences contact stability, modulation of tether abundance selectively tunes the lateral growth and integrated area of contact sites without substantially altering contact-site density. By combining inducible expression with chemically or optically triggered dimerization, our platform enables continuous, predictable control over ER-PM contact-site size and total cellular contact area. This dual-parameter strategy, independent control of intermembrane spacing and tether density, provides a versatile framework for dissecting how the abundance and spatial organization of membrane tethers govern the structural and functional outputs of ER-PM contact sites.

## CONCLUSIONS

We have developed two complementary and independently tunable platforms for precise, reversible, and quantitative control of ER-PM contact sites, combined with high-contrast visualization in living cells. The ABA-inducible CID system provides nontoxic, reversible chemical control with dose-dependent regulation of contact-site formation and disassembly kinetics, while the iLID/SspB optogenetic system enables rapid, reversible, light-dependent modulation with superior temporal resolution ; both allow dose-dependent regulation of contact-site formation kinetics and abundance, manifested as increased contact-site density and total area fraction per cell, without substantially changing the mean size of individual contact sites. In contrast, systematic variation of α-helical linker length or inducible tether expression precisely tunes the lateral growth and stability of individual contact sites, reflected in increased mean size, total contact area, and fluorescence intensity, without altering their density. These complementary mechanisms establish predictive structure–function relationships for membrane contact sites and define independent parameters to modulate contact-site dynamics, abundance, geometry, and stability.

Beyond providing versatile experimental tools, this dual-parameter framework has broad implications for dissecting how contact-site architecture influences essential cellular processes, including Ca^2+^ signaling, lipid transport, metabolic coordination, and directional migration. The quantitative and reversible nature of these systems allows for precise, perturbation-free studies of ER-PM contact site dynamics, enabling repeated induction-recovery cycles and systematic exploration of spatial and temporal regulation. Importantly, the modular design of these platforms is generalizable, providing a blueprint for manipulating other inter-organelle contact sites and expanding the experimental toolkit for studying intracellular organization, signaling compartmentalization, and disease-relevant dysregulation. Altogether, our work establishes a versatile, predictive, and high-fidelity framework for engineering and interrogating membrane contact sites, bridging molecular design to cellular function with unprecedented precision.

## AUTHOR INFORMATION

### Author Contributions

R.Z. conceived the project. S.Z., J.F., and R.Z. designed experiments, interpreted the data and wrote the manuscript. S.Z., Y.Z., and J.F. performed experiments. S.Z. analyzed the data. R.Z. acquired funding and supervised the project.

## ACKNOWLEDGMENT

This work was supported by National Institute of General Medical Sciences (R35GM142973 to R.Z.)

## Supporting Information

**Figure S1.**
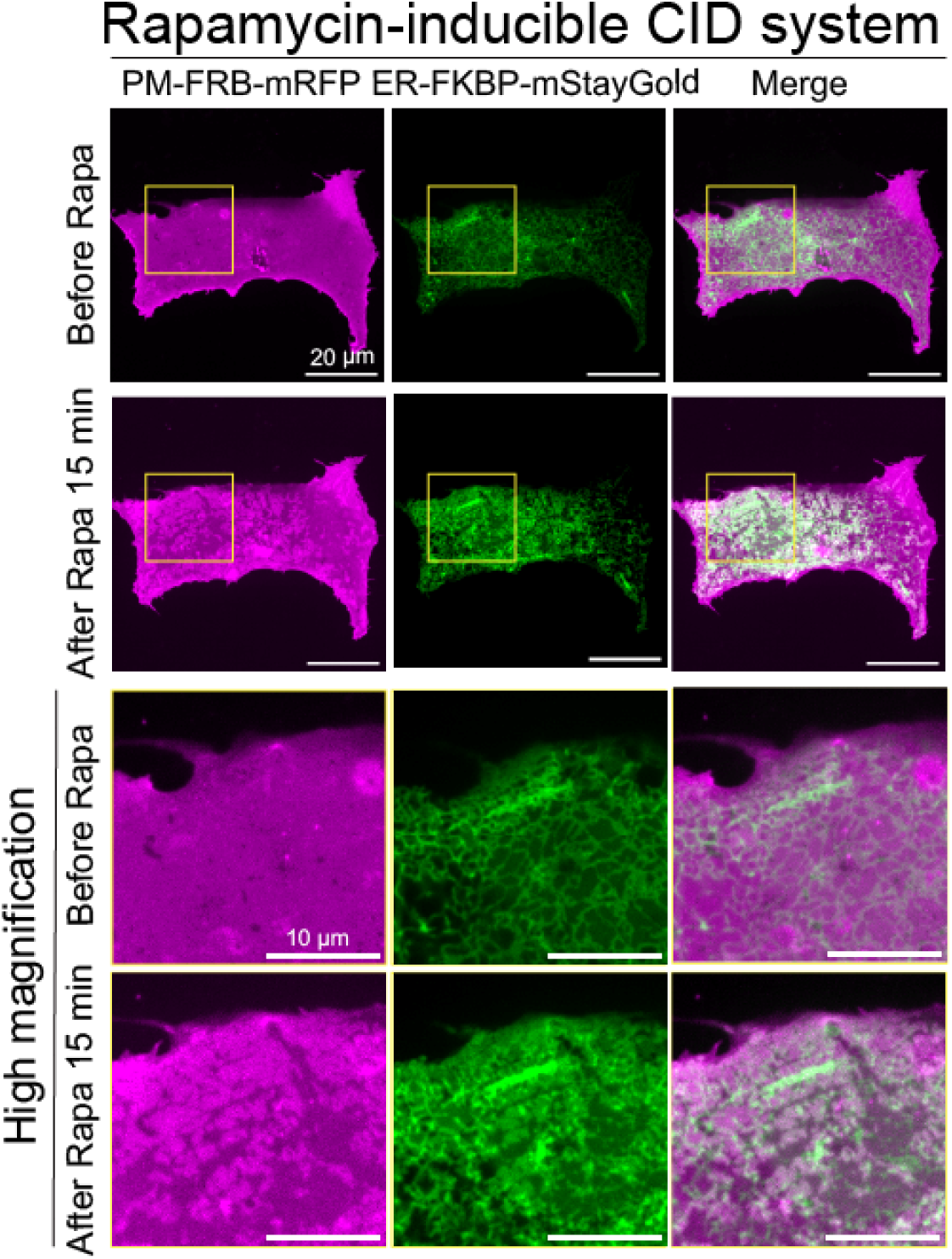
Representative confocal images of U2OS cells expressing the rapamycin-inducible construct (PM-FRB-mRFP-T2A-mStayGold-FKBP-ER) before and 15 min after treatment with 100 nM Rapamycin. Yellow boxed regions are shown at higher magnification below. Scale bar: 20 μm (original view); 10 μm (magnified view).

**Figure S2.**
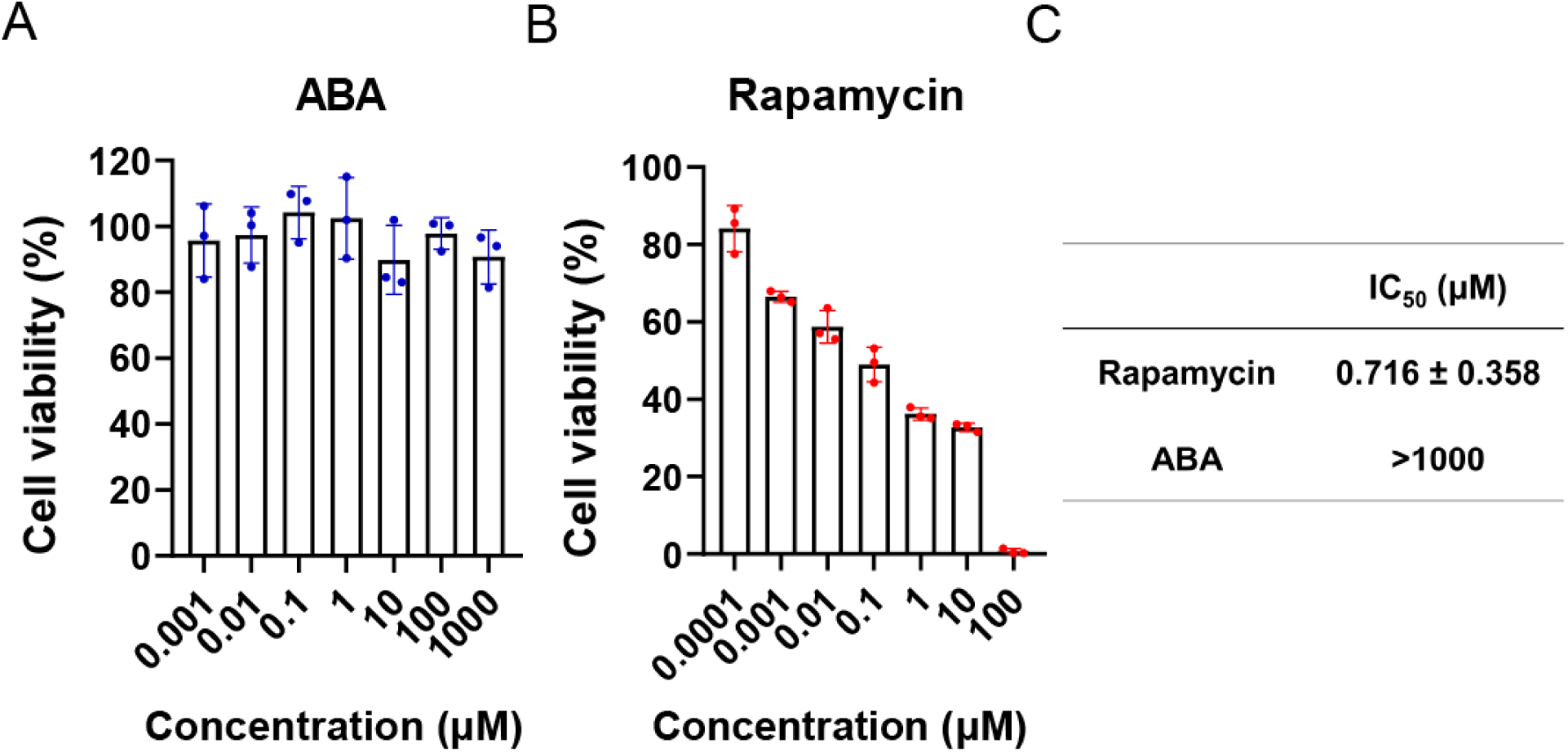
(**A, B**) MTT assay of U2OS cells treated with increasing concentrations of ABA (**A**) or rapamycin (**B**). Cell viability is expressed as relative viability (% of untreated control). Data represent mean ± s.e.m. from three biological replicates (*n* = 3). (**C**) IC values of rapamycin and ABA in U2OS cells, determined by nonlinear regression analysis of dose-response curves using GraphPad Prism 9. Data represent mean ± s.e.m. from three independent biological replicates (*n* = 3).

**Figure S3.**
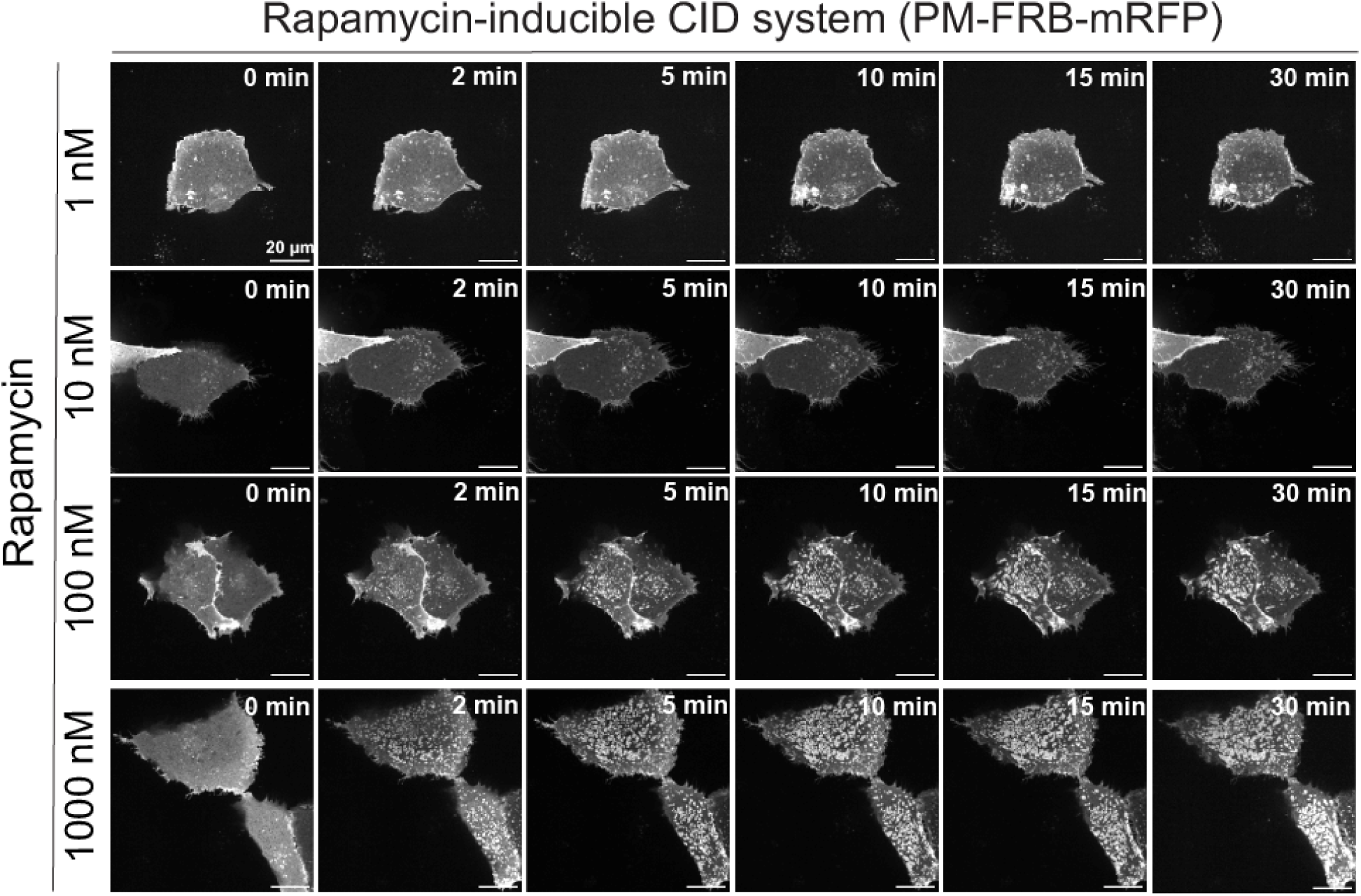
(**E**) Representative time-course (0-30 min) and dose-response images of rapamycin-induced ER-PM contact site formation in U2OS cells. Cells were transfected with rapamycin-inducible CID construct and imaged by confocal microscopy 48 h after transfection. Scale bar: 20 μm.

**Figure S4.**
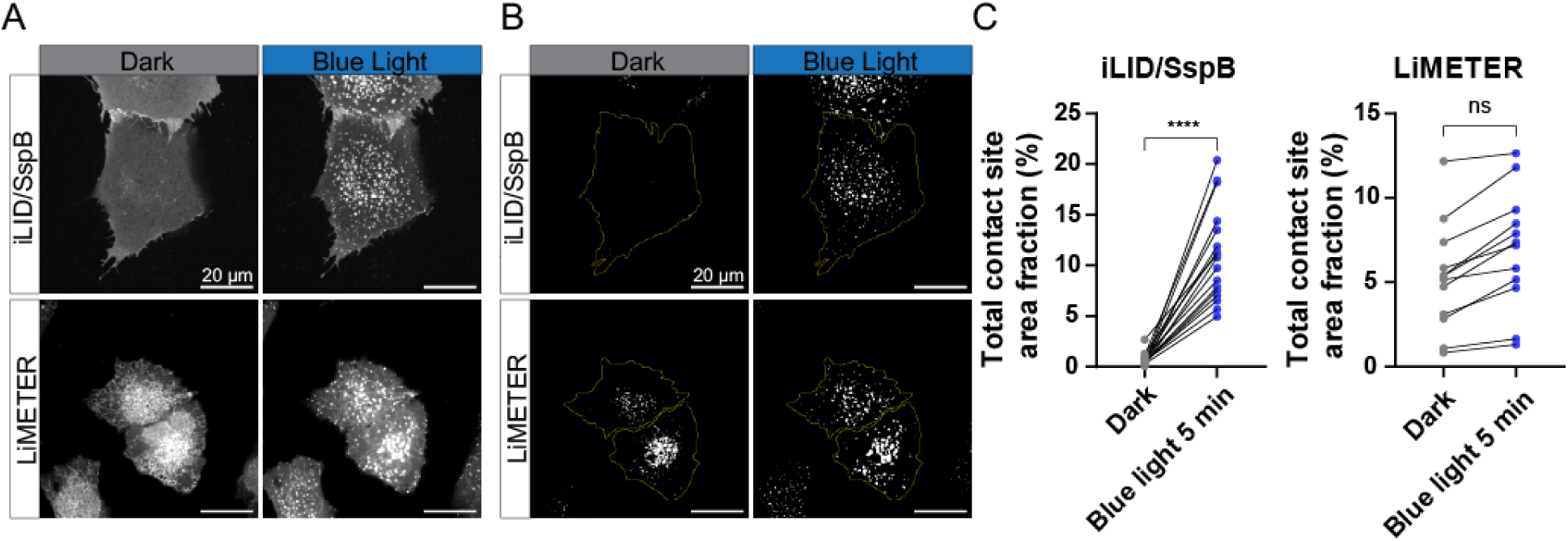
(**A**) Representative confocal images of the ventral surface of U2OS cells expressing iLID/SspB LID construct (mRFP channel) or LiMETER construct (GFP channel) before (left) and 5 min after (right) blue-light illumination. A 470-nm laser (∼142 mW/cm^2^, measured at the sample plane) was used to activate the iLID/SspB or LiMETER system, and GFP signals were simultaneously acquired for LiMETER. Scale bar: 20c μm. (**B**) Segmentation masks of the ventral surface of U2OS cells shown in Fig. S4A, generated by MATLAB-based image analysis to identify punctate ER-PM contact sites. These masks were used as the cell area for contact site quantification. (**C**) Quantification of total ER-PM contact site area fraction before and after blue-light illumination in U2OS cells expressing iLID/SspB (*n* = 17 cells) or LiMETER (*n* = 12 cells) construct, based on the ventral surface confocal images shown in Fig. S4A. P values were calculated using two-tailed unpaired Student’s *t* tests*: *P < 0.05, **P < 0.01, ***P < 0.001, ****P < 0.0001*, and ns, not-significant.

**Figure S5.**
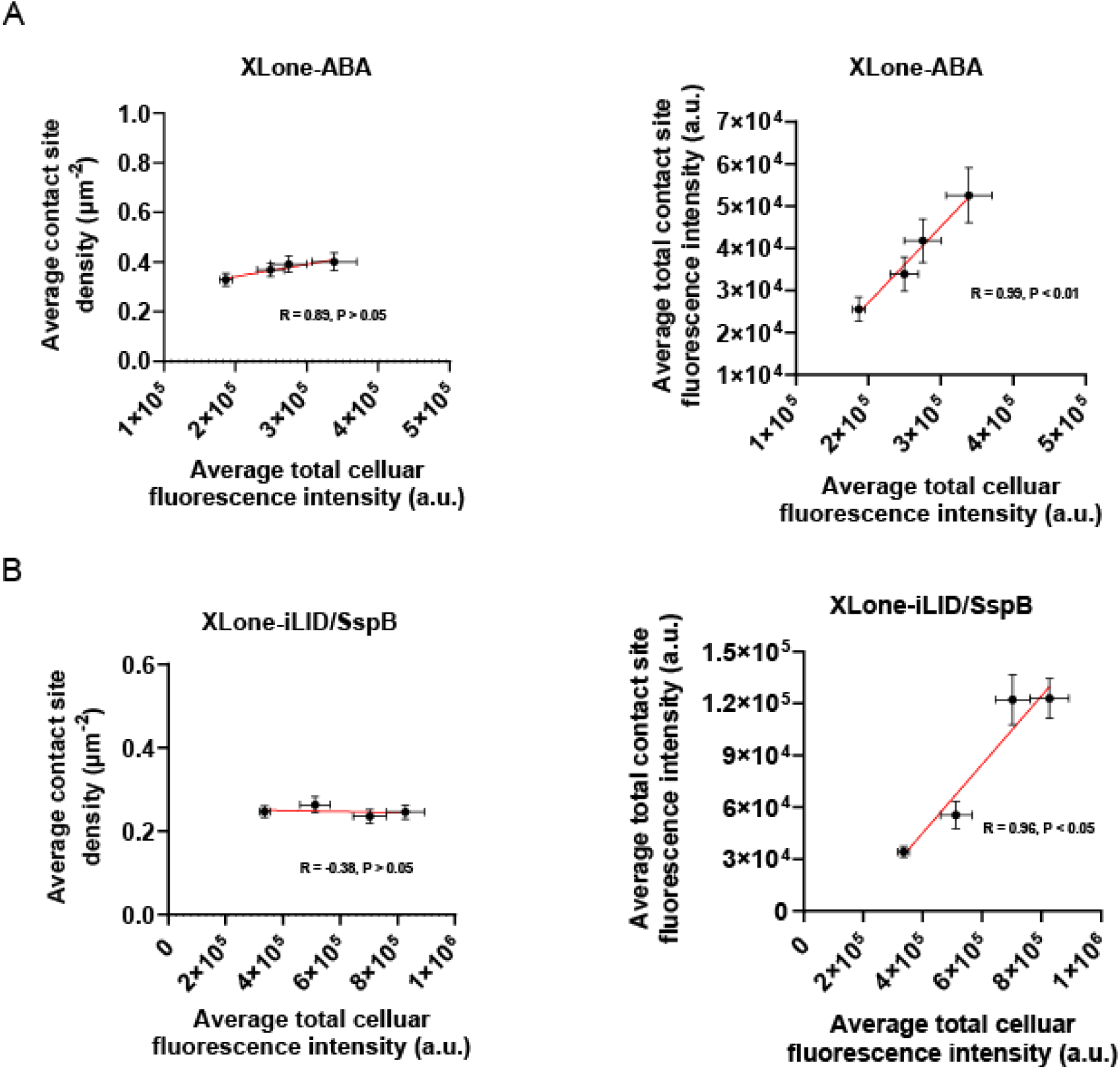
Correlations between average total cellular fluorescence intensity and ER–-PM contact site parameters, including average contact site density (left) and average total contact site fluorescence intensity (right) in XLone-ABA-inducible CID (**A**) or XLone-iLID/SspB LID (**B**) systems. Pearson correlation coefficients (R) and corresponding P values were calculated using GraphPad Prism 9.

